# Enrichment analysis for spatial and single-cell metabolomics accounting for molecular ambiguity

**DOI:** 10.1101/2024.08.23.609355

**Authors:** Bishoy Wadie, Martijn R. Molenaar, Lucas Maciel Vieira, Theodore Alexandrov

## Abstract

**Summary:** Imaging mass spectrometry (imaging MS) has advanced spatial and single-cell metabolomics, but the reliance on MS1 data complicates the accurate identification of molecular structures, not being able to resolve isomeric and isobar molecules. This prevents application of conventional methods for overrepresentation analysis (ORA) and metabolite set enrichment analysis (MSEA). To address this, we introduce *S2IsoMEr* R package and a web app for METASPACE, which uses bootstrapping to propagate isomeric/isobaric ambiguities into the enrichment analysis. We demonstrate *S2IsoMEr* for single-cell metabolomics and the METASPACE web app for spatial metabolomics.

**Availability and Implementation:** METASPACE web app can be used on existing and new datasets submitted to METASPACE (https://metaspace2020.eu). The source code for the *S2IsoMEr* R package is available on GitHub (https://github.com/alexandrovteam/S2IsoMEr).

## Introduction

Recent advances in imaging mass spectrometry (imaging MS) have boosted the emerging fields of spatial metabolomics and lipidomics as well as opened novel avenues for obtaining metabolomics data on the single-cell level. Due to the limitations on sample amounts and design of modern imaging mass spectrometers, imaging MS data is usually collected in the so-called MS1 mode, without the untargeted MS/MS tandem fragmentation commonly used in bulk metabolomics. Molecular annotation of MS1 data provides identities on the Level 2 of the Metabolomics Standards Initiative (Sumner *et al*., 2007), mainly relying on the m/z values of ions and their isotopic peaks (Palmer *et al*., 2017). As a result, molecular candidates with identical (‘isomers’) or similar (‘isobars’) mass-to-charge (m/z) values cannot be resolved.

This unavoidable ambiguity in the identification of metabolites complicates downstream analyses such as overrepresentation analysis (ORA) and metabolite set enrichment analysis (MSEA) (Xia and Wishart, 2010b, 2010a; Picart-Armada *et al*., 2018). In these types of analyses, metabolites are linked to shared properties such as metabolic pathway association or molecular class, after which the enrichment of these properties within the dataset is determined using statistical methods (Zhao and Rhee, 2023). A common way to handle the ambiguous identification in imaging MS is to manually curate the candidates and select the most plausible ones. However, this does not always remove the molecular ambiguity, introduces a selection bias and is time-consuming, especially when the number of metabolites is large.

Various metabolite enrichment methods are available, including overrepresentation-based, ranking-based, and network topology-based approaches (Zhao and Rhee, 2023). Popular metabolomic data analysis platforms such as MetaboAnalyst (Pang *et al*., 2024) offer robust enrichment methods utilising diverse resources and network-based algorithms to aid in interpretation. However, these methods are primarily designed for bulk and LC-MS/MS-based metabolomics. They do not address the molecular ambiguity of annotations in imaging MS studies and are not yet adapted for spatial and single-cell metabolomics.

Here, we present two implementations of a new method to perform metabolite enrichment analysis that addresses the challenge of metabolite identification ambiguity. The first is a web app on METASPACE (Palmer *et al*., 2017). The second, *S2IsoMEr*, is a new R package designed for (spatial) single-cell metabolomics datasets. In both implementations, the key idea to handle molecular isomers and/or isobars is to propagate the molecular ambiguity into the analysis by iterative random sampling (hereafter, bootstrapping) of isomeric/isobaric candidates.

## Description

METASPACE is an online platform for metabolite identification in spatial metabolomics. It allows users to upload imaging MS datasets and to perform annotation of metabolites and lipids in a false discovery rate (FDR)-controlled manner (Palmer *et al*., 2017). In the annotation process, METASPACE makes use of databases containing molecular formulas of biological interest. For each molecular formula, METASPACE calculates a score based on features such as isotopic and spatial distribution, based on which a corresponding false discovery rate (FDR)-value is assigned. Because METASPACE performs annotation based on the MS1 data, it reports multiple possible isomers and isobars for the annotated ions which is mainly determined by the chosen annotation database (e.g. HMDB). For the ion [C_42_H_82_NO_8_P+H]^+^, for example, it reports the isomeric lipids PE 37:1 and PC 34:1. As a result, each annotated ion may be associated with a large number of molecular structures.

Due to this annotation ambiguity, performing enrichment analysis of molecule names using a classical approach is not possible. Accordingly, we leveraged bootstrapping to consider such ambiguity in the enrichment analysis. In each iteration, one molecular candidate for each ion is randomly sampled (with replacement) from all isomeric and/or isobaric candidates corresponding to that ion (**Figure 1A**), optionally using weights indicating their likelihood or relative abundance. The bootstrapping procedure is repeated for *m* iterations. Afterwards, aggregate statistics of the individual enrichment analyses are calculated and reported (**Figure 1A**). More details on the methods are available in **Supplementary Note 1**.

**Figure 1.**
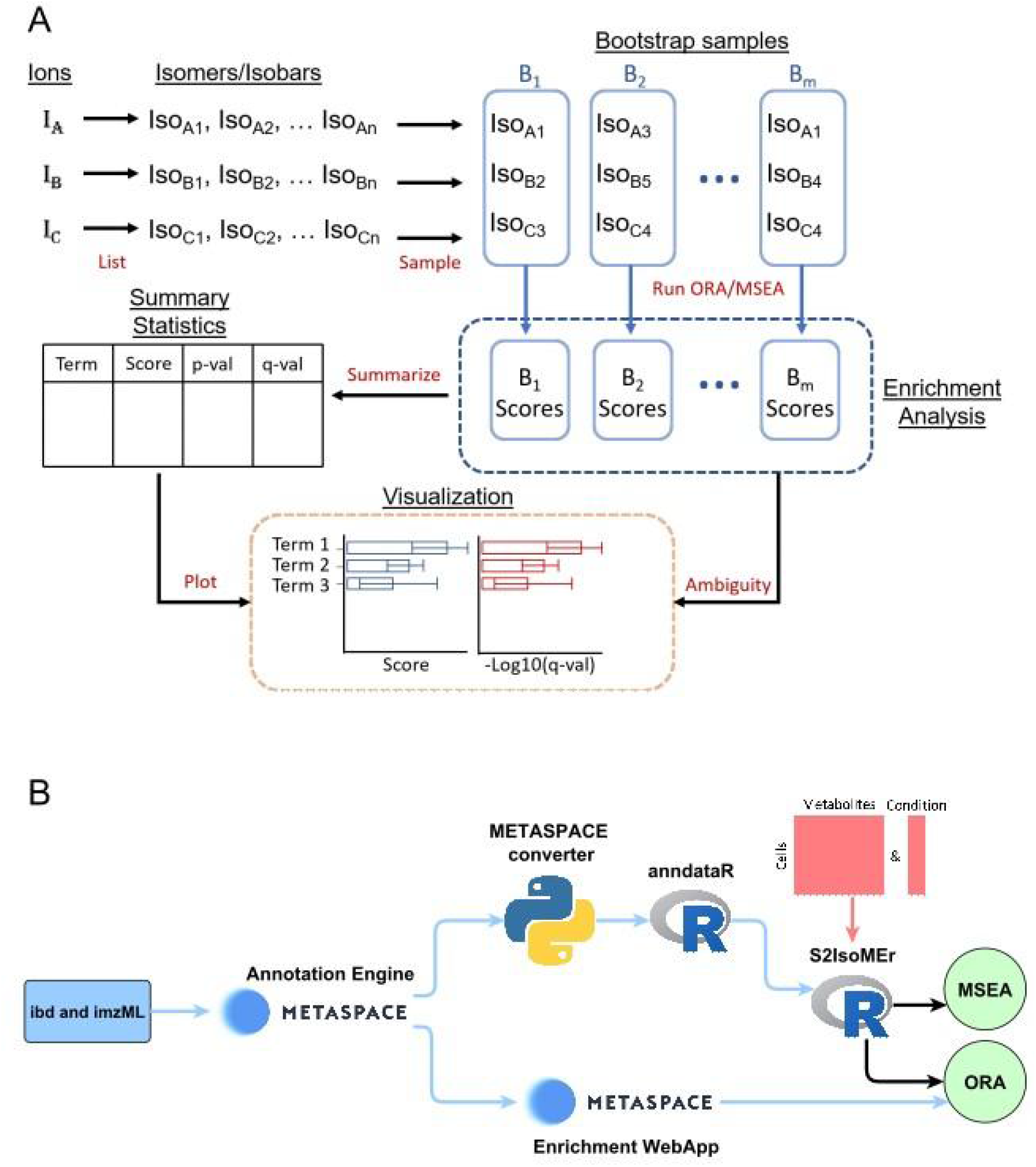
Overview of Bootstrapping-based enrichment and implementations. (A) Illustration of molecular ambiguity handling with bootstrapping and result reporting. (B) Flowchart of input for the enrichment web app and *S2IsoMEr*, with blue and red edges representing spatial and single-cell metabolomics workflows, respectively.

Regarding the property databases (hereafter metabolite sets) required for enrichment analyses, the algorithm supports LION ontology for lipids (Molenaar *et al*., 2019) and multiple class-based and pathway-based sets for metabolites and lipids curated from RAMP-DB (Braisted *et al*., 2023) (**Supplementary Note 2)**. This database encompasses biological pathways and metabolic classes, which are further categorized into super, main, and sub-classes. The pathways from RAMP-DB integrate multiple resources, including SMPDB, Reactome, KEGG, and WikiPathways (Frolkis *et al*., 2010; Fabregat *et al*., 2018; Kanehisa and Goto, 2000; Kutmon *et al*., 2016)

In the METASPACE web app implementation for ORA, the user selects a metabolite set when submitting a dataset. Once processing has finished, users can access the enrichment web app and view the results. Bootstrapping is performed as mentioned previously on the chosen set and enrichment of each term is assessed against its presence in the full molecular database in each bootstrap sample using a one-tailed Fisher’s exact test. The resulting *p*-values from the Fisher’s exact test are adjusted for multiple testing using the Benjamini and Hochberg method (Benjamini and Hochberg, 1995) and median intersection size along with fold enrichment score are reported per term (**Supplementary Note 1.4**). Bar plots present the aggregated statistics from the hypergeometric tests across bootstraps, and users can adjust significance thresholds, FDR cutoffs, and explore the enriched annotations for each term (**Figure S1**).

Although Imaging MS is primarily known for spatial metabolomics, it has also been increasingly adopted for single-cell metabolomics as well (Rappez *et al*., 2021; Capolupo *et al*., 2022). The resulting datasets are matrices consisting of *n* metabolites measured for *m* single cells. While the METASPACE implementation performs ORA of metabolite annotations from one dataset (as compared to all metabolites from the database used for annotation), *S2IsoMEr* is currently designed for single-cell metabolomics and supports both ORA and MSEA (**Figure 1B**). It makes use of the same metabolite sets available in the METASPACE web app and can be run with or without accounting for isomeric/isobaric ambiguity. The decision tree in **Figure S2** provides guidance on how to select the appropriate enrichment type. Moreover *S2IsoMEr* can be extended to support spatial metabolomics datasets from METASPACE using *metaspace-converter (https://github.com/metaspace2020/metaspace-converter)*. This tool converts metaspace datasets to an anndata object and can subsequently be loaded into R with the *anndataR* R package (Virshup *et al*., 2023, 2021) to provide single-pixel matrices for *S2IsoMEr* (**Figure 1B**). *S2IsoMEr* requires as inputs (i) a matrix containing the single-cell metabolomics measurements, (ii) a vector indicating the condition or group to which each cell belongs, and (iii) the conditions of interest to compare (*e*.*g*., for comparing two conditions or cell types). Under the hood, *S2IsoMEr* associates the provided molecular formulas to isomeric and, optionally, isobaric molecular structures, based on the internal molecular databases also used in METASPACE. It is assumed that cells from both groups share the same metabolite annotations but differ in their intensities. Therefore, the user has the option to perform MSEA by ranking all metabolites prior to enrichment. In each bootstrap iteration, all metabolites are ranked using a selected local statistic such as *t*-values from the Welch Two Sample *t*-test, BWS, Wilcoxon or Log Fold changes (Zyla *et al*., 2017) (**Supplementary Note 1.3**). For each metabolite set (*e*.*g*. a LION term or a metabolite set provided by a user), an enrichment score is calculated using either the Kolmogorov–Smirnov (KS) statistic or the normalized enrichment score (NES) implemented in the *fgsea (Korotkevich et al*., *2021*) R package.

## Case studies

### Bootstrapping-based overrepresentation analysis of spatial metabolomics dataset in METASPACE

We showcase ORA in METASPACE using a mouse brain dataset (https://metaspace2020.eu/dataset/2022-05-31_10h46m34s). For the enrichment analysis, we selected the CoreMetabolome database (Wadie *et al*., 2024) as an annotation database with an FDR cutoff of 10%, excluded off-sample annotations as implemented in METASPACE and performed ORA using LION ontology as metabolite set (settings: Database=CoreMetabolome, molecule type=lipid, category=class, ontology=LION).

To visualize the results, the implemented METASPACE web app provides three main views: a table, a chart, and data filtering options. The table view displays all enrichment values associated with the current dataset, along with relevant data used for enrichment calculation. Users can export this data as a CSV file for further analysis. The chart view presents enrichment results as a bar chart, providing essential information and interactivity. By clicking on an ontology name or bar, users are redirected to the annotation page, where they can explore potential molecules involved in enrichment annotation (**Figure S3**). Lastly, the data filtering feature allows users to refine results using criteria such as ontology database, FDR, annotation database, off-sample, and p-value threshold. These filters dynamically update the table and chart views to reflect the selected parameters.

As anticipated, LION terms associated with sphingolipids such as ‘phosphosphingolipids’ and ‘sphingolipids’ were found to be highly overrepresented (**Figure S1**) consistent with previous reports that brain tissues have relatively high amounts of sphingolipids (Hussain *et al*., 2019; Olsen and Færgeman, 2017).

### Metabolite set enrichment analysis of single-cell metabolomics data with the S2IsoMEr R package

To showcase *S2IsoMEr*, we applied it to our previously reported single-cell metabolomics dataset (Rappez *et al*., 2021) of HepaRG cells, which models NASH by stimulating HepaRG cells with fatty acids and other inhibitors compared to a healthy control, followed by MALDI imaging MS..In total, there were 3 perturbation conditions and 1 control condition, each with 4 replicates, resulting in 16 datasets in METASPACE (**Supplementary Table 1**) that cover the 4 conditions. The single-cell matrices per condition were concatenated in a single matrix and only annotations reported in METASPACE were considered. More details on samples are available in the corresponding MetaboLights record (https://www.ebi.ac.uk/metabolights/MTBLS78).

Both ORA and MSEA analyses were conducted using the combined dataset and compared against metabolite sub-classes curated from RAMP-DB, as previously mentioned, with HMDB v4.0 (Wishart *et al*., 2018) serving as the annotation database. Since METASPACE provides FDR-controlled annotations, we selected annotations with a 10% FDR threshold as the query set and included all reported annotations as the universe for enrichment analysis. For ORA, annotations were split into upregulated and downregulated markers based on the sign of log fold changes (LFC) between the cells of the specified conditions..

We compared HepaRG cells treated with exogenous fatty acids (‘F’, representing a steatosis model) to untreated cells (‘U’, reflecting a healthy state). We performed a bootstrapping metabolite enrichment analysis with 100 iterations, accounting for isomeric and isobaric structures. As anticipated and consistent with previous analyses, terms associated with fat accumulation, such as ‘triacylglycerols’ and ‘diradylglycerols,’ were highly enriched in steatotic cells, while ‘glycerophosphocholines’ and ‘glycerophosphoethanolamines’ were enriched in healthy cells, in both ORA and MSEA (**Figures S4, S5**). The findings were also consistent when the other metabolic states were compared to healthy cells (**Figure S6**).

## Conclusion

Ambiguous identification of metabolites and lipids in spatial and single-cell metabolomics by imaging MS complicates downstream enrichment analysis approaches that are routinely performed on bulk metabolomics data. In this work, we described a method of bootstrapping enrichment analysis to address this problem by performing the analysis multiple times, with iterative random sampling of the molecular candidates belonging to each ion. This approach circumvents the laborious and error-prone manual selection of plausible molecules and acknowledges the intrinsic annotation uncertainty. The implementation of bootstrapping-based enrichment in METASPACE provides an intuitive and accessible way to quickly explore the coverage of lipid and metabolite classes and properties in imaging MS datasets submitted to METASPACE. In addition, the *S2IsoMEr* R package extends the bootstrapping-approach to ranking-based enrichment in single-cell metabolomics datasets and we foresee a wide adoption of the package as more single-cell datasets are generated.

## Supplementary data

The single-cell metabolomics dataset used to showcase *S2IsoMEr* was previously published (Rappez *et al*., 2021) and is available at the MetaboLights repository under the accession number MTBLS78 (https://www.ebi.ac.uk/metabolights/MTBLS78).

## Conflict of interest

T.A. holds patents in spatial and single-cell metabolomics and is a co-founder of a startup in single-cell metabolomics at the BioInnovation Institute.

## Funding

We acknowledge the funding from the European Research Council (Consolidator grant No. 773089), European Horizon2020 grants NEARDATA and CloudSkin (grant No. 101092644, 101092646), Swiss National Science Foundation project PROMETEX, Michael J Fox Foundation, La Caixa Banking foundation under the project code HR23-00516, and Chan-Zuckerberg Initiative (CZI).

## Notes

https://www.ebi.ac.uk/metabolights/MTBLS78

https://metaspace2020.eu/dataset/2022-05-31_10h46m34s

https://github.com/bisho2122/bmetenrichr

