## Supplementary Information for "Enrichment analysis for spatial and single-cell metabolomics accounting for molecular ambiguity"

### Supplementary Note 1 :

#### 1. Bootstrapping

To generate a bootstrapped sample from a list of possible isomeric/isobaric molecules, we employed a resampling strategy. For each of the *N* bootstrapping iterations, the process involves sampling 1 molecule from the list of isomers/isobars for each given ion in the annotation list. It is assumed that all isomers/isobars are equally likely and have the same sampling probability. However, annotation weights can be provided in *S2IsoMEr a-priori*.

Given:

- *M* : Number of input ions
- $Iso_i$ : Set of isomers/isobars (molecules) associated with ion *i*, where $i \in \{1, 2, \ldots, M\}$ and the size of $Iso_i$ may vary for each ion.
- $w_i$ *:* The vector of sampling probabilities (weights) corresponding to the isomers in $Iso_i$

For each ion $i \in \{1, 2, \ldots, M\}$, a single molecule $x_i$ is sampled from $Iso_i$. This process is repeated for *N* bootstrapping iterations.

The bootstrapping process can be expressed as:

$$B^{(n)} = \left\{x_1^{(n)}, x_2^{(n)}, \ldots, x_M^{(n)}\right\} for\, n = 1, 2, \ldots N \quad (1)$$

Where:

- $x_i^{(n)}$ is the isomer sampled from $Iso_i$ during the *n-th* iteration.
- $x_i^{(n)} \sim Sample\left(Iso_i, w_i\right)$, where "Sample" represents a weighted random selection using $w_i$ as the probabilities.

#### 2. ORA contingency table adjustment

Overrepresentation analysis (ORA) in *S2IsoMEr* is performed using re-implementation of the "run_ora" function from the decoupleR package (Badia-I-Mompel et al. 2022). Everything is performed as standard ORA with a one-tailed fisher exact test. Additionally, since each observed ion typically corresponds to only a single isomer in the bootstrapping process, whereas multiple isomers of the same molecular formula may exist in the expected molecules for a given term in the chosen metabolite set, special handling was applied. Specifically, false negatives were filtered to exclude the remaining isomers for a given formula if the sampled isomer is a true positive (i.e. mapped to the given term in the metabolite set) to avoid inflating the count of false negatives due to redundant representations of the same molecule.

#### 3. MSEA signed ranking

In MSEA, metabolites are ranked based on their intensity changes between the conditions under comparison. In *S2IsoMEr* we implemented 4 ranking metrics : Log Fold Change (LFC), Wilcoxon rank-sum statistic, t-test statistic and BWS (Zyla et al. 2017). LFC measures the ratio of intensity levels between two conditions on a logarithmic scale, indicating how much a metabolite abundance increases or decreases. Positive LFC values show higher abundance in the test condition, while negative values indicate lower abundance. The Wilcoxon rank-sum is a non-parametric method that ranks all metabolites from both conditions and compares the sum of ranks to assess differences without assuming normal distribution. On the other hand, t-test is a parametric test that compares the means of two conditions under the assumption of normality. We use the statistic value exclusively for running MSEA, and to improve result interpretation, we multiply these statistical values by the sign of the Log Fold Change (LFC). This approach ensures that the direction of regulation (up or down) is incorporated while maintaining the magnitude of the statistic.

a) LFC :

$$mean_{i,x} = \frac{1}{|C_x|} \sum_{j \in C_x} Sc_{ij} \quad (2)$$

$$mean_{i,y} = \frac{1}{|C_y|} \sum_{j \in C_y} Sc_{ij} \quad (3)$$

$$LFC_i = \log_2 (mean_{i,y}) - \log_2 (mean_{i,x}) \quad (4)$$

Where $C_x$ and $C_y$ are the sets of cell indices for the reference and test conditions, respectively. $Sc_{ij}$ represent the log10 intensity level of metabolite $i$ in cell $j$ . $mean_{i,x}$ and $mean_{i,y}$ are the mean of log10 intensity of metabolite $i$ in reference and test conditions, respectively.

b) Wilcoxon rank-sum statistic

$$wilcox.stat_j = \text{sign}(LFC_j) \cdot \sum_{i=1}^{n_x} r_i - \frac{n_x (n_x + 1)}{2} \quad (5)$$

Where $n_x$ is the number of cells in test condition *x*, $r_i$ denotes the rank of the *i-th* cell in both reference and test conditions for metabolite $j$, and $LFC_j$ is LFC of metabolite $j$.

c) BWS

$$B(N,n,R) = \frac{1}{n}\sum_{i=1}^{n}\frac{\left(R_{i-1} - i\cdot\frac{N}{n}\right)\cdot\left|R_{i-1} - i\cdot\frac{N}{n}\right|}{\frac{i}{n+1}\cdot\left(1-\frac{i}{n+1}\right)\cdot\left((N-n)\cdot\frac{N}{n}\right)} \quad (6)$$

$$BWS.stat_j = \frac{1}{2}\left(B\left(N, n_{x_j}, R_{x_j}\right) - B\left(N, n_{y_j}, R_{y_j}\right)\right) \quad (7)$$

The above equation represents the modified one-tailed (greater) Neuhauser's difference statistic denoted by Murkami as a modification of the Baumgartner-Weiss-Schindle (BWS) statistic (Murakami 2012).

- $x_j$ : Intensity of metabolite $j$ in cells of test condition
- $y_j$ : Intensity of metabolite $j$ in cells of Reference condition
- $n_{x_j}$ : Number of cells in test condition
- $n_{y_j}$ : Number of cells in Reference condition
- $R_{x_j}$ : Rank of $x_j$ relative to $y_j$
- $R_{y_j}$ : Rank of $y_j$ relative to $x_j$
- $N : n_{x_j} + n_{y_j}$

#### 4. Enrichment scores

In MSEA, the reported enrichment score is the Normalized Enrichment Score (NES) in the fgsea (fast gene set enrichment analysis) method (Korotkevich et al. 2021). NES is a metric that adjusts the raw enrichment score (ES) to account for differences in metabolite set sizes. In MSEA, the enrichment score indicates how well a set of metabolites is represented at the extremes (top or bottom) of a ranked list of metabolites. The NES is calculated by normalizing the ES to a distribution of scores obtained from random permutations of the metabolite sets. For ORA, we use the fold enrichment (FE) score which measures how much a specific metabolite set or pathway is overrepresented compared to what would be expected by chance. It is calculated as the ratio of the proportion of observed metabolites in a given pathway (or set) (*geneRatio)* to the proportion of all metabolites that are mapped to a given pathway (or set) (*BgRatio)* (Wu et al. 2021). The formula for the Fold Enrichment is

$$FE = \frac{observed}{\text{expected}},\ observed = \frac{TP}{TP + FP},\ \text{expected} = \frac{TP + FN}{TP + FP + FN + TN} \quad (8)$$

Where TP, FP, FN, TN denote true positives, false positives, false negatives and true negatives respectively.

5. Isomeric ambiguity score

To quantify the isomeric/isobaric ambiguity for each ion we used Shannon's entropy of weighted probabilities for molecular isomers associated with a particular ion. The ambiguity score quantifies how uncertain or ambiguous the assignment of a single ion is when multiple isomers are possible. The entropy is computed based on the distribution of weights assigned to its isomers. The entropy is 0 if there is only one possible isomer and for multiple isomers (N>1), the score increases as the distribution of weights becomes more uniform, indicating higher uncertainty.

$$Ambiguity = -\sum_{i=1}^{N_j} p_i \cdot \log_2(p_i) \quad (9)$$

Where $p_i$ represents the probability of each isomer/isobar and $N_j$ is the total number of possible isomers/isobars for ion $j$.

When the probabilities are equally weighted, each isomer would have the same probability. If there are N isomers, the probability for each isomer is $p_i = 1/N_j$.

#### Supplementary Note 2 :

The metabolite sets provided for both the METASPACE web app and S2IsoMEr were curated from either the LION ontology for lipids (Molenaar et al. 2019) or from RAMP-DB for metabolite- and lipid-specific classes and pathways (Braisted et al. 2023). The metabolite sets were constructed using the SQL database dump of RAMP-DB (v2.2.1). For metabolite classification sets, HMDB served as the source database, with only records labelled as quantified or detected being selected. The selected classification types were main, sub, or superclass categories based on the ClassyFire classification system (Djoumbou Feunang et al. 2016). For each class, two sets were created—one mapping terms to molecular formulas and another mapping terms to molecule names, using chemical information from the "chem_props" table in the database. For lipid classes, the same process was followed, but with LipidMaps as the source instead of HMDB. Additionally, biological pathways were mapped to lipids and metabolites using information from the "pathway" and "analytehaspathway" tables in the SQL database. For more details on the available metabolite sets, refer to the documentation for the "Load_background" function in *S2IsoMEr* (https://alexandrovteam.github.io/S2IsoMEr/reference/Load_background.html) .

### Supplementary Figures:

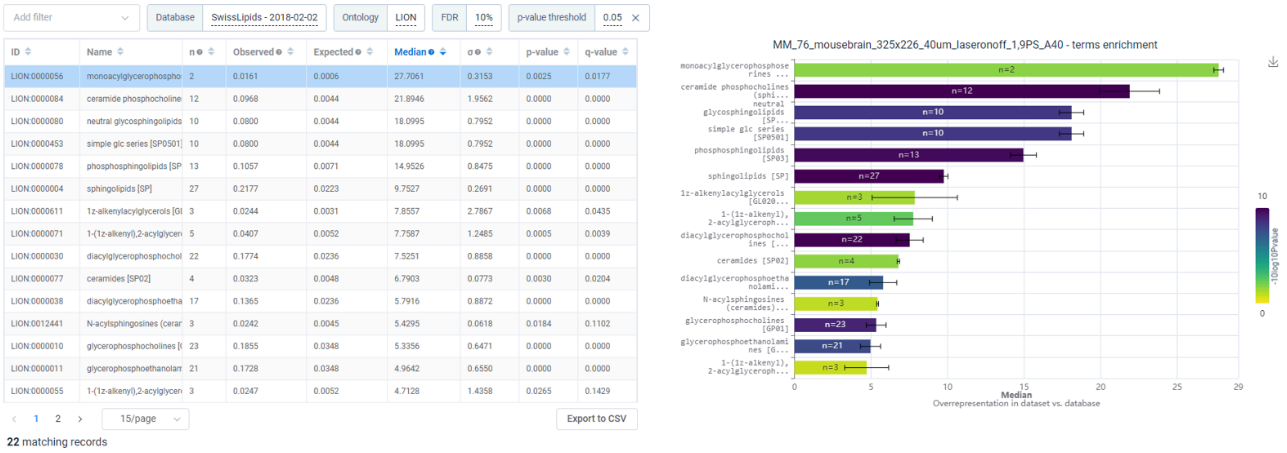

| ID | Name | n | Observed | Expected | Median | σ | p-value | q-value |
|---|---|---|---|---|---|---|---|---|
| LION:0000056 | monoacylglycerophospho | 2 | 0.0161 | 0.0006 | 27.7061 | 0.3153 | 0.0025 | 0.0177 |
| LION:0000084 | ceramide phosphocholine | 12 | 0.0968 | 0.0044 | 21.8946 | 1.9562 | 0.0000 | 0.0000 |
| LION:0000080 | neutral glycosphingolipids | 10 | 0.0800 | 0.0044 | 18.0995 | 0.7952 | 0.0000 | 0.0000 |
| LION:0000453 | simple glc series [SP0501] | 10 | 0.0800 | 0.0044 | 18.0995 | 0.7952 | 0.0000 | 0.0000 |
| LION:0000078 | phosphosphingolipids [SP | 13 | 0.1057 | 0.0071 | 14.9526 | 0.8475 | 0.0000 | 0.0000 |
| LION:0000004 | sphingolipids [SP] | 27 | 0.2177 | 0.0223 | 9.7527 | 0.2691 | 0.0000 | 0.0000 |
| LION:0000611 | 1z-alkenylacylglycerols [Gl | 3 | 0.0244 | 0.0031 | 7.8557 | 2.7867 | 0.0068 | 0.0435 |
| LION:0000071 | 1-(1z-alkenyl),2-acylglycer | 5 | 0.0407 | 0.0052 | 7.7587 | 1.2485 | 0.0005 | 0.0039 |
| LION:0000030 | diacylglycerophosphochol | 22 | 0.1774 | 0.0236 | 7.5251 | 0.8858 | 0.0000 | 0.0000 |
| LION:0000077 | ceramides [SP02] | 4 | 0.0323 | 0.0048 | 6.7903 | 0.0773 | 0.0030 | 0.0204 |
| LION:0000038 | diacylglycerophosphoetha | 17 | 0.1365 | 0.0236 | 5.7916 | 0.8872 | 0.0000 | 0.0000 |
| LION:0012441 | N-acylsphingosines (cerar | 3 | 0.0242 | 0.0045 | 5.4295 | 0.0618 | 0.0184 | 0.1102 |
| LION:0000010 | glycerophosphocholines [( | 23 | 0.1855 | 0.0348 | 5.3356 | 0.6471 | 0.0000 | 0.0000 |
| LION:0000011 | glycerophosphoethanolam | 21 | 0.1728 | 0.0348 | 4.9642 | 0.6550 | 0.0000 | 0.0000 |
| LION:0000055 | 1-(1z-alkenyl),2-acylglycer | 3 | 0.0247 | 0.0052 | 4.7128 | 1.4358 | 0.0265 | 0.1429 |



*Figure S1 : METASPACE Enrichment web app*
Screenshot from METASPACE web app enrichment results on a brain datasets ([https://metaspace2020.eu/dataset/2022-05-31_10h46m34s](https://metaspace2020.eu/dataset/2022-05-31_10h46m34s)) using FDR 10%, a p-value threshold of 0.05, LION ontology as metabolite set and SwissLipids as annotation database.

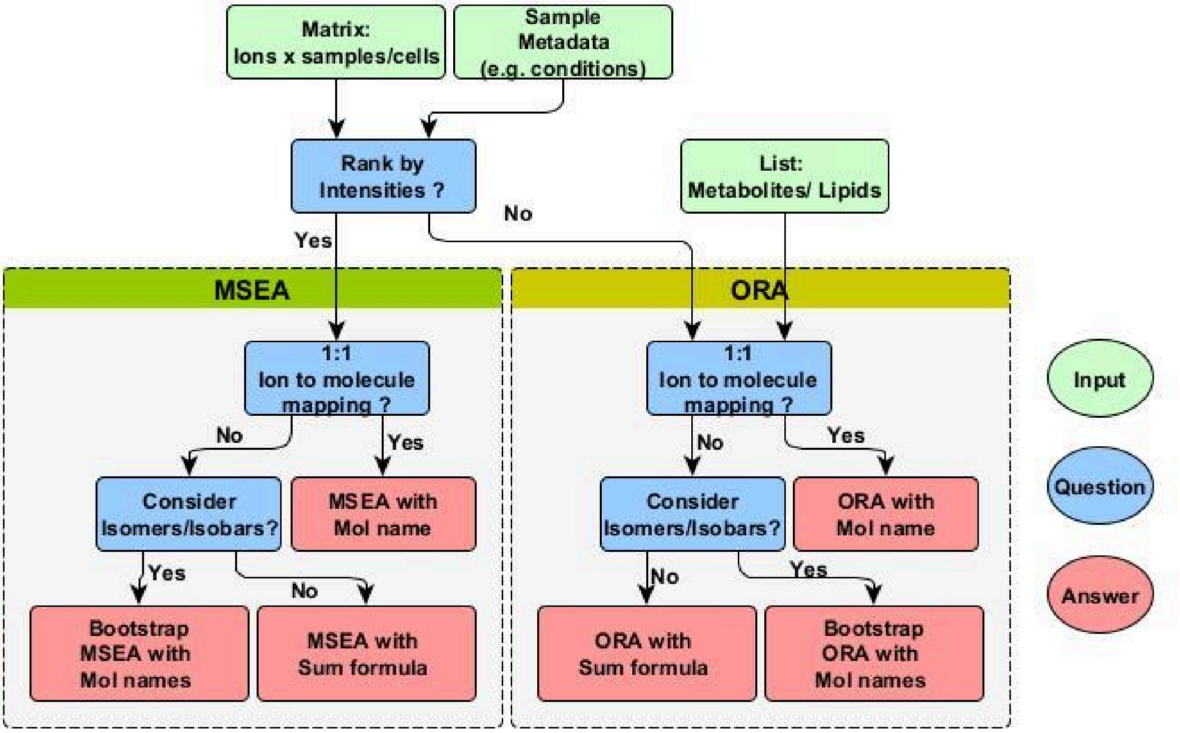



*Figure S2 : Enrichment Types Decision Tree*
Flowchart to help the user decide on which enrichment type would be appropriate based on the provided input data. Nodes coloured in green, blue and red represent input, questions and answers, respectively.

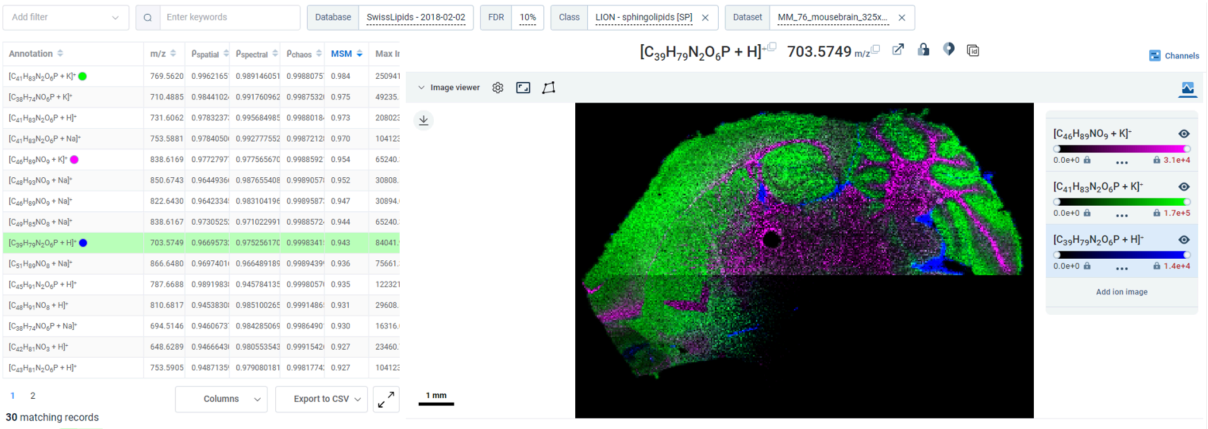


*Figure S3 : Sphingolipids annotations in mouse brain*
Screenshot from annotation page on METASPACE for brain dataset (https://metaspace2020.eu/dataset/2022-05-31_10h46m34s) after clicking on sphingolipids bar in *Figure S1.*

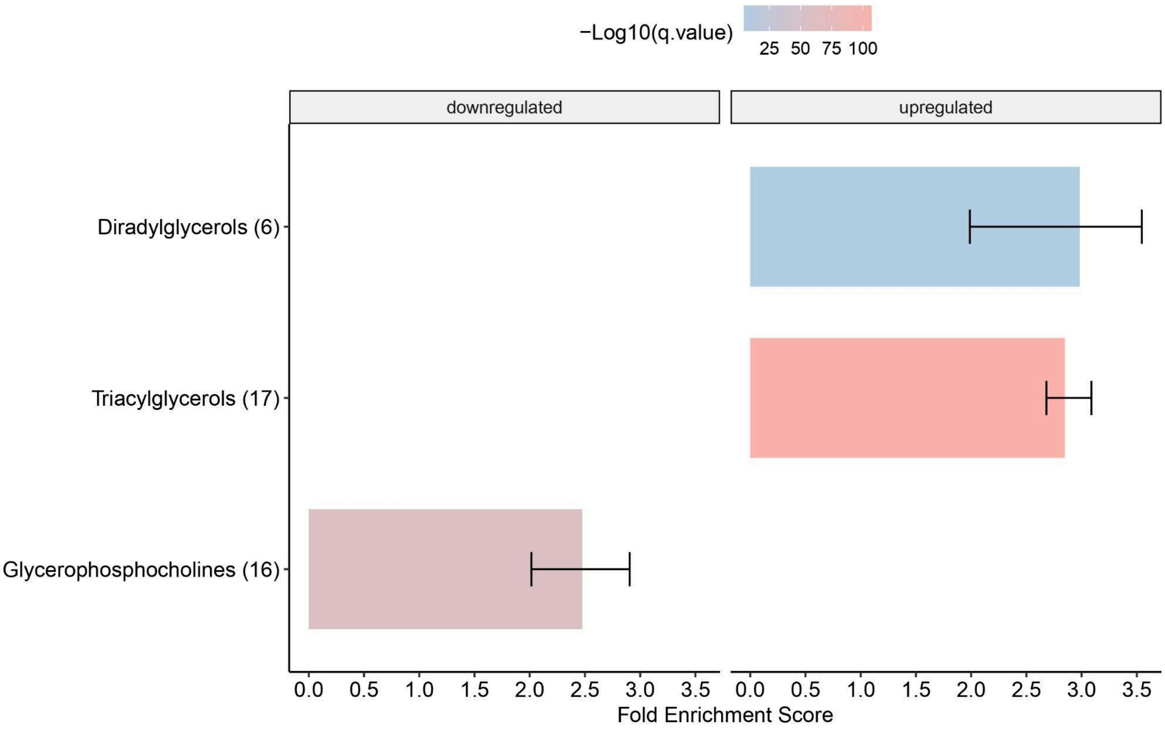


*Figure S4 : ORA results for SpaceM NASH dataset*
Barplot showing results of bootstrapping-based ORA on a single-cell dataset based on a NASH model from SpaceM (https://www.ebi.ac.uk/metabolights/MTBLS78). Enriched terms are plotted on the y-axis and fold enrichment score on the x-axis. Bar colour corresponds to -log10 (q-value) and error bars represent min and max enrichment scores across bootstrap iterations. Size of term/query overlap (i.e. number of molecules) are displayed in parentheses next to enriched terms.

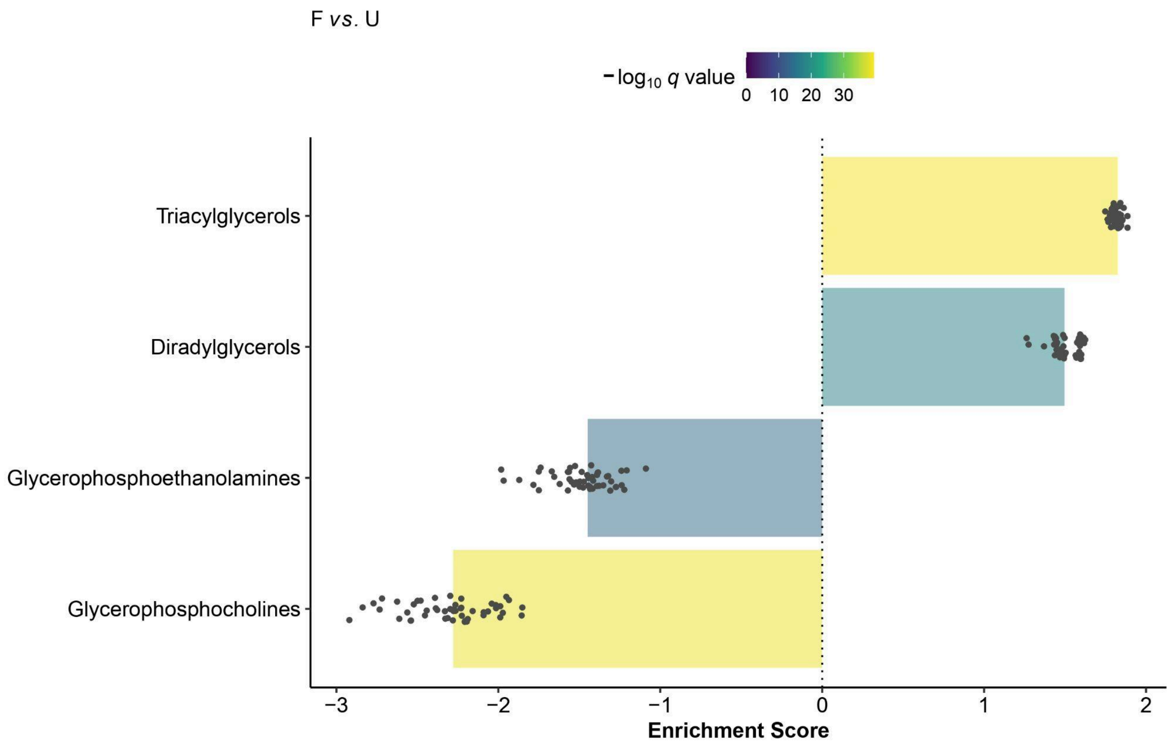


*Figure S5 : MSEA results for SpaceM NASH dataset*

Barplot showing results of bootstrapping-based MSEA on a single-cell dataset based on a NASH model from SpaceM (https://www.ebi.ac.uk/metabolights/MTBLS78). Enriched terms are plotted on the y-axis and normalized enrichmed score (NES) on the x-axis. Bar colour corresponds to -log10 (q-value) and jittered points represent NES scores for each bootstrap iteration.

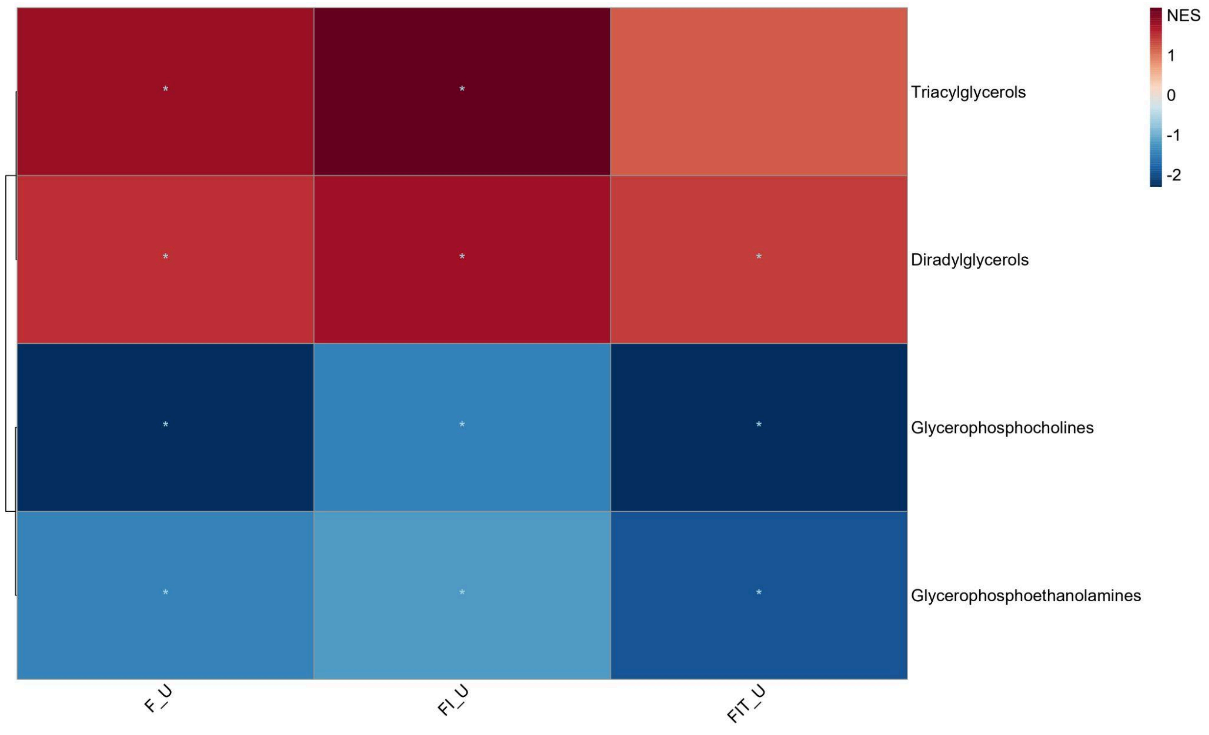

*Figure S6 : MSEA Multi-condition results*
Heatmap showing MSEA results of bootstrapping-based MSEA on a single-cell dataset based on a NASH model from SpaceM (https://www.ebi.ac.uk/metabolights/MTBLS78). Rows represent the enriched terms and columns representing pairwise conditions as “test_reference”. Color scale corresponds to normalized enrichment score (NES).
